# Snooping helices : The elastic path finding algorithm of growing hyphae

**DOI:** 10.1101/2025.11.12.685335

**Authors:** Antoine Rittaud, Elodie Couttenier, Sophie Bachellier-Bassi, Christophe d’Enfert, Igor M. Kulić, Catherine Villard

**Author notes:** co-first authors (Antoine Rittaud, Elodie Couttenier). These two authors contributed equally to this work (Igor M. Kulić, Catherine Villard). Author Contributions: A.R. performed all mathematical modeling and associated data analysis and image segmentation. E. C. performed all experiments and analyzed some of the presented data (Fig. 1) under the supervision of S. BB. and C. d’E. IM K. and A.R. set up the mathematical model and conceptual framework of data analysis. All authors discussed the results and contributed to the final manuscript. C. V. and IM K. supervised the project and the findings of this work. Competing Interest Statement: The authors declare no competing interests.

## Abstract

How living organisms utilize physical mechanisms to sense their environments and make informed decisions is an open question at the interface of biology and physics. In filamentous organisms like fungal hyphae, the decisions are taken by their growing tip cells and later imprinted onto the rest of the multi-cellular filament. Here we report on the growth and pathfinding of hyphae from the opportunistic fungal pathogen *Candida albicans*, whose ability to cross intestinal epithelial layers is associated to severe systemic infections in humans. It has been sporadically reported that *C. albicans*’s hyphae display helical growth inside or on top of agar gels, helicity turning in the latter case into two-dimensional oscillatory shapes. We provide an extended description of oscillatory *C. albicans* hyphal growth modalities, revealed under various physical confinements thanks to the use of dedicated microfluidic devices and quantitative time-lapse imaging-based analysis. These include sudden sliding events accompanied by curvature switching of the tip portion, resulting in a final oscillatory morphology of the entire filament, and stable curved tips moving against vertical microfluidic channel’s walls. These behaviors are unified under the formalism of growing squeezed helices, in which the final hyphal curved shapes result from an elastic energy minimization of a spatially confined helical portion at the tip followed by a continuous solidification front. Ultimately, the combination of our experimental results and theoretical framework provide an insight into the penetration strategy of *C. albicans* hyphae, which is essential for the virulence of this fungal microorganism.

**Significance Statement:** Proprioception is the integrated sense of self-movement and body position in complex organisms. Here we describe a novel, mechanical form of proprioception driving directional choice making in tip-growing helical organisms. We show that *C. albicans* hyphae utilize their built-in helicity as an environment-scanning mechanism to explore their surrounding and find target surfaces for invasion. When confined to surfaces, hyphae continue producing in-plane oscillatory shapes that promote further invasive behavior. *C. albicans’* inherent mechanical instabilities regulate the switching of growth direction and their abrupt directional decisions can be understood as elastic bifurcations of squeezed, confined helices.

## Introduction

Helically coiled filaments are ubiquitous in Nature. They are observed at different scales, ranging from molecular structures such as intermediate filaments [1], microtubules [2], flagella [[3]], bacteria [4] up to multicellular structures such as climbing vines and roots [[5]. The helical growth of the plant root, named circumnutation by Charles Darwin, has recently been identified as crucial for the exploration and penetration of soil-mimicking substrates [6], which gives this generic geometry of life high biological and ecological relevance. This strategy used by plants is also a source of inspiration in the field of robotics, stimulating the development of motion planning algorithms to improve the performance of soft robots [7]. Helical growth is a widely shared behavior outside plants in the tree of life. Early cell divisions can occur in an helical arrangement in a variety of animals, leading to the well-known coiling of the shell in gastropods or the twisted body of the roller mutants of the nematode *Caenorhabditis elegans* (see [8] for a review).

In Fungi, which share turgor pressure-driven growth with plants, oscillatory growth modalities of hyphae were reported sporadically and coined as diversely as coils [9], zigzags [10], kinks [11], sinusoids or helices [12, 13]. The three last cited studies were performed on the human opportunistic fungal pathogen *Candida albicans*, a benign member of the human microbiota that can turn into one of the deadliest human opportunistic fungal pathogens. *C. albicans* is a polymorphic fungus that replicates either as a budding yeast or as a ≈ 2µm-diameter filament (hypha) composed of mono-nucleated compartments. Among the reversible *C. albicans* switching systems (yeast-to-hypha but also white-opaque [14]) occurring in response to environmental conditions [15], the yeast-to-hypha switch is recognized as a key virulence trait of this important human fungal pathogen [16]. Interestingly, *C. albicans* helical hyphae have been reported during active human onychomycosis, indicating that they may play a role in invasion [17]. They also have been observed *in vitro* in agar-invasion assays, and associated with increased virulence in mice models [11] or increased resistance to antifungal agents [18], suggesting a possible mechanistic link between helicity and pathogenicity.

Here we propose a unified mechanistic description of *C. albicans* hyphal oscillatory growth modalities observed under various conditions, implying either adhesion or physical confinement which we fine-tune using microfluidic technologies. Both are instrumental in the invasive lifestyle of *C. albicans*, as its hyphae may adhere and navigate at the surface of epithelial cells (drawing oscillations, see [19]) or experience confinement in e.g. the crypts of the colon intestine [20].

We show that the repertoire of hyphal growth dynamics and morphology is well captured by the extension of the concept of “squeelix” (squeezed helix), previously formulated by us and co-workers [21], to growing *C. albicans* hyphae under confinement. In the present model, we consider that a hypha is composed of a frozen (resignated) portion growing with time and an elastically reconfigurable (mechanically frustrated) portion close to the tip. A dynamic frontier sets the boundary conditions for the frustrated part, whose shape at every growth step results from the minimization of the total elastic energy stored in it. Last but not least our approach captures generic behaviors that promote efficient penetration mechanisms, suggesting that the intrinsic helicity of *C. albicans* hyphae could be the result of a selection process that favors its virulence.

## Results

### *C. albicans* hyphae display oscillatory behaviors under physical confinement

We first incorporated *C. albicans* yeasts (SC5314 reference strain) within a mixture of filamentation-inducing medium and different concentrations of agar, from 0.2 to 5%. The sample is then cast on a thin PDMS hydrophilic layer where it solidifies, producing gels of various elastic moduli in the range of 100-2000 kPa [22], and put in the incubator at 37°C and 5% CO_2_. We observed hyphae growing in all directions from the yeasts embedded in the bulk of the gel, reaching both the upper (with air) and lower (with PDMS) interfaces. In all cases, we noticed oscillations: hyphae display helical shapes within the gel, meandering morphologies at the agar-air interface, and in-plane sinusoidal-like oscillations in-between the gel and the PDMS hydrophilic surface (**Fig.1A**). Although right-handed helices only have been reported in the quasi-2D situation represented by a cellophane sheet on top of agar [18], we observed both right- and left-handed helical hyphae growing in the volume of agar gels (**Figure S1 and Supp. Movie 1**). In the interface configurations (i.e. agar-air and PDMS-agar), *C. albicans* filaments experience confinement. Due to the presence of a thin water layer, confinement of growing hyphae at the interface agar-air is mediated by the normal forces resulting from water surface tension directed toward the agar surface (this confinement can be destroyed by adding water on top of the gel, preventing curved shapes in favor of straight hyphae). In-plane confinement also rules the growth of hyphae at the PDMS-agar interface. In this case where friction can be easily tuned by the rigidity of agar, we observed an increased proportion of oscillating hyphae, characterized by higher oscillation amplitudes, with increased agar concentration (**Figure S2**). Note that hyphae do not oscillate on hydrophilic PDMS or glass covered with liquid medium devoid of agar. Lastly, transferring hyphae from such substrates, which are not conducive to cell adhesion (and thus from where they can be easily detached), to a cell culture treated Petri dish restores oscillatory growth similar to that observed in confinement (**Supp. Movie 2**).

**Figure 1:**
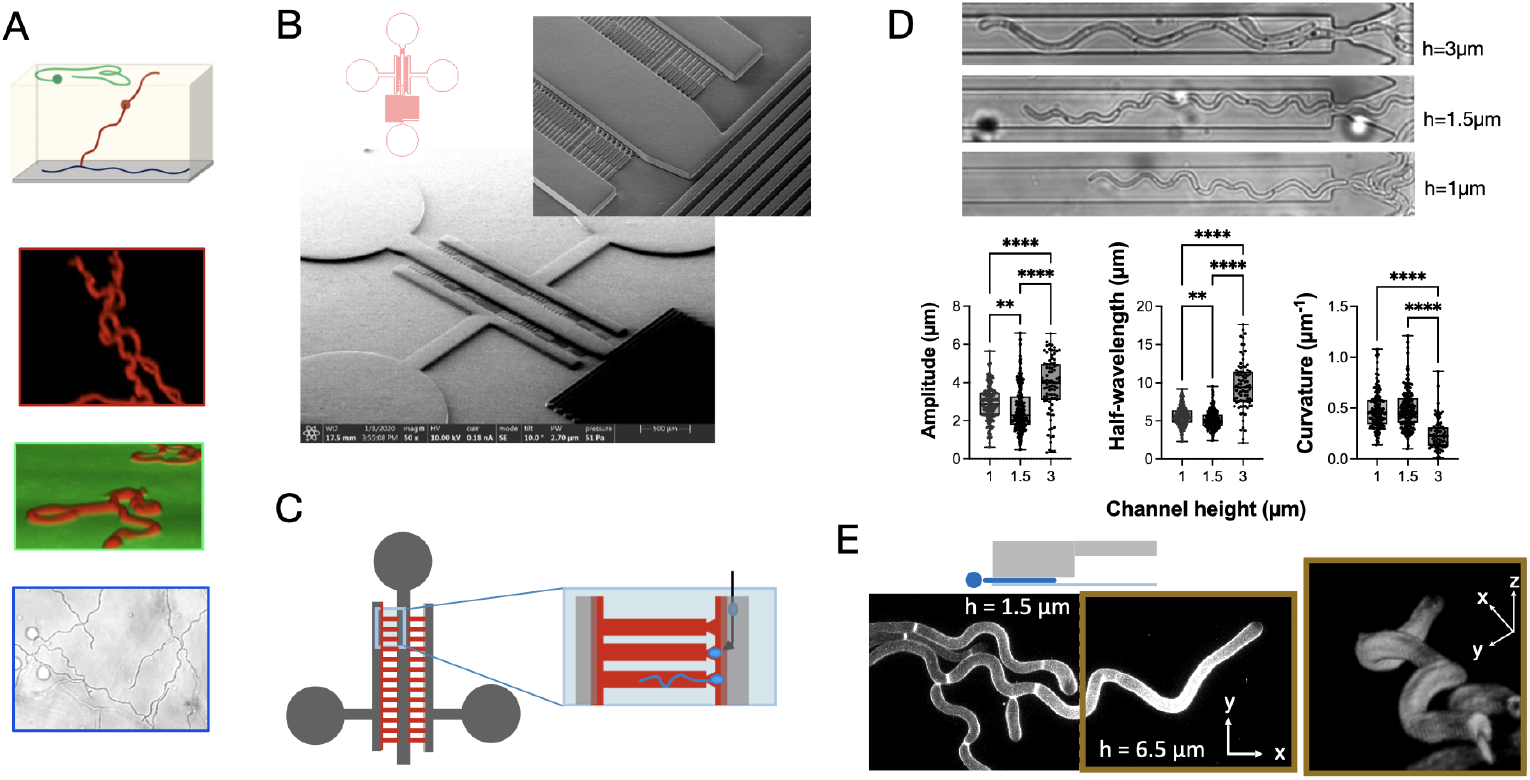
**A -** Scheme of hypha locations in yeasts-seeded agar gels: agar-air interface (green), agar bulk (red) and PDMS-agar interface (blue). The outlines of the images illustrating these situations use the same color code. Top image: calcofluor white (CW) staining (red) of an hyphae growing in a 2% agar gel. Middle image: the surface of the gel is stained with Dextran (green) and the hyphae with CW (red). Bottom image: phase contrast image of hyphae navigating in-between a 5% agar gel and an hydrophilic PDMS layer. Hypha’s approximate diameter: 2µm. **B -** Scheme and scanning electron microscopy of the mold of the devices used in this study, comprising 4 inlet-outlets, a central channel terminated by a serpentine and opening on two mirror arrays of the microchannel. The zoomed-in image shows two-height microchannels. **C -** Scheme describing the principle of seeding and chip operation. **D -** Phase contrast images of hyphae in microchannels of different heights. Graph: distribution of amplitudes, half-wavelengths and maximum curvatures (whose expression is 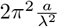, with *a* the amplitude and *λ* the oscillation wavelength) characterizing oscillations as a function of channel height for a channel width of 10µm. (n_hyphae_, n_oscillations_)= (17,176) for h=1µm, (19,223) for h=1.5µm and (11,83) for h=3µm. **E -** Scheme of bi-height channel cross-section. Confocal imaging: projected view of CW stained hyphae and 3D view of the 6.5µm high channel portion (brown square outlines). **Fig.S1** - Curvature and torsion of *C. albicans* helical growth in agar cells. **Movie 1**: Stack of 3D reconstruction in 2% agar gel. **Fig.S2** - *C. albicans* oscillations at the interface PDMS-agar as a function of agar concentration. **Movie 2**: Timelapse of glass to Petri dish. **Movie 3a and 3b**: Two examples of confocal 3D stack of hyphae in bi-heigth 10µm wide (Movie 3a) and 40µm wide (Movie 3b) channels.

In view of these preliminary results, we decided to turn to microfluidic devices to fine-tune physical confinement. We devised microfluidic channels of h=1, 1.5 and 3µm height and of various width (from *W* =1.5 to 40µm). Considering the 2µm mean hypha diameter, the smallest channel heights should promote strong to moderate squeezing, while the highest height would only ensure loose confinement. In a more sophisticated type of device, channels display alternative heights of 1.5 and 6.5µm in order to successively confine and partly de-confine a same hypha (**Fig. 1B**). Briefly (see Methods for more details), yeasts are seeded in a central channel through a main inlet. A portion of these cells are diverted toward trapping pockets located at the entrance of microchannels where they quickly start to initiate a germinal tube which will elongate into a long hypha (**Fig. 1C**).

Hyphae observed in phase-contrast imaging oscillate markedly in microfluidic channels independently of their height (see images of **Fig. 1D**). However, the mean amplitude, wavelength and maximum curvature of these oscillations allow us to distinguish between the squeezed condition from the one where hyphae are only loosely maintained in height. For instance, the maximum curvature is not statistically different between h=1 and 1.5µm and h=3µm (see graphs of **Fig. 1D**). We then turned to bi-height microchannels combining moderate confinement and significant de-confinement (see the zoom image of Fig. 1B) in order to follow along a same hypha the effect of a change in z-confinement. We observed in phase contrast imaging a striking difference between hyphal morphologies in each channel portion, the de-confined hypha displaying loose meandering (**Supp. Movie 3a and 3b**). When resolved in 3D using confocal imaging (see an example in **Fig. 1E**), the same projected image reveals a truncated helix, changing, in the particular case shown in Fig. 1E, its direction of rotation after interacting with the PDMS ceiling of the chip, which may prove too low for the helix to grow unhindered. These results led us to consider in the following mainly fairly squeezed hyphae for an in-depth study of the static and dynamical oscillatory hyphal shapes.

### Statistical properties of hyphae: Environmental “snooping” and long range persistence

To quantify the statistical behavior of oscillating confined hyphae, we borrowed methods from polymer physics of semiflexible filaments, with the most intuitive observable being their tangent-tangent-correlation function, TTCF(Δ*s*), as a function of the arc-length distance Δ*s*. For random, spatially uncorrelated Gaussian-distributed curvature variation along the length, one would expect a purely exponential decay of the tangent correlation - as found in thermally fluctuating semi-flexible “wormlike chains” (WLC) [23]. However, both in 3D agar gels and on 2D surfaces at the PDMS/agar interface the hyphae robustly displayed a characteristic behavior that is quite different from predictions of the WLC model, see **Fig.2**.

**Figure 2:**
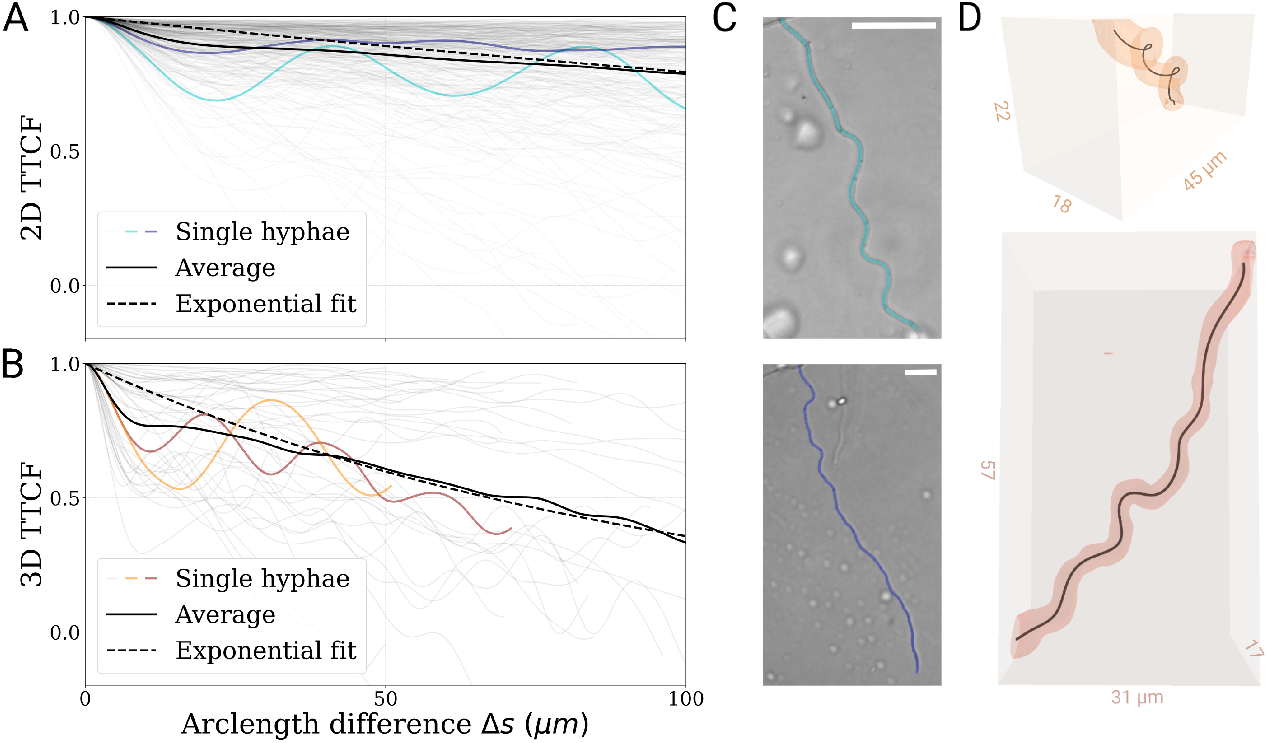
Directional persistence of hyphae in agar gels. **A -** Tangent-tangent correlation (TTCF) functions of hyphae center lines at a PDMS-agar interface, two corresponding hyphae are displayed in **C** with their centerlines traced. The blue-colored hypha oscillates slightly, and maintains its global orientation. The cyan one displays clear oscillatory behavior while also maintaining its global orientation. The average correlation function (black) is fitted with an exponential decay (dotted black) of characteristic decay length 433µm. **B -** TTCF of hyphae center lines in agar gel volume, two corresponding hyphae are represented in **D**. The orange hypha displays helical behavior, while the red one displays an oscillatory behavior taking effect in a virtual plane within the 3D gel. The exponential decay fit in 3D shows a characteristic decay length of 97µm. **Movie 4a and 4b**: 3D rotational view of the helical (Movie 4a) and planar (Movie 4b) hyphae shown in **Fig.2D**.

At short distances the TTCF falls off quicker than linear, resulting in a quadratic drop, so that the correlations at this scale are strongly reduced. However, after this initial drop, the TTCF recovers, crossing over to the typical exponential decay, as expected within the WLC model. The 2D and 3D situations differ by both their exponential decay characteristic length and the range of deviation from this exponential decay. Both are coherent with the fact that an additional dimension can only impair directional persistence. This combined short and long distance environment exploratory can be characterized as “persistent snooping”. The short distance oscillatory snooping acts as a deterministic scanning process with a built-in typical wavelength and seems to be utilized to efficiently probe the environment while allowing for a continuation of the initial direction if no better conditions are encountered on the short scale.

### Rapid rearrangements of hypha tip portions generate oscillations

In the following we asked ourselves the question of how the oscillations are dynamically formed in microchannels. The most immediate idea is that the newly emerging apex would continuously draw the oscillations as it moves forward, while the rest of the hyphae remains shape invariant. However, dynamic monitoring of hyphal growth under confinement shows that this is not the case. **Fig.3** (see also **Supp. Movie 5a and 5b**) indicates that, unexpectedly, oscillations are formed by successive sliding of a limited portion of the hypha close to the apex, while the rest of the hypha upstream stays still. In other terms, a growing hypha seems to be constituted of two parts: one reconfigurable below the apex, and a second one, whose length increases continuously, showing an invariant shape that can be described as “solidified” or “frozen”. We estimated the length of the reconfigurable part by computing the displacement in-between sliding events along the hypha as a function of the distance to the tip (see **Fig.S3**). An exponential fit of the mean displacement gives a characteristic decay length of 5.9µm with a threshold of measurable displacement around a length of 25µm, which is interestingly close to the typical length of a mononucleated hypha compartment [24].

**Figure 3:**
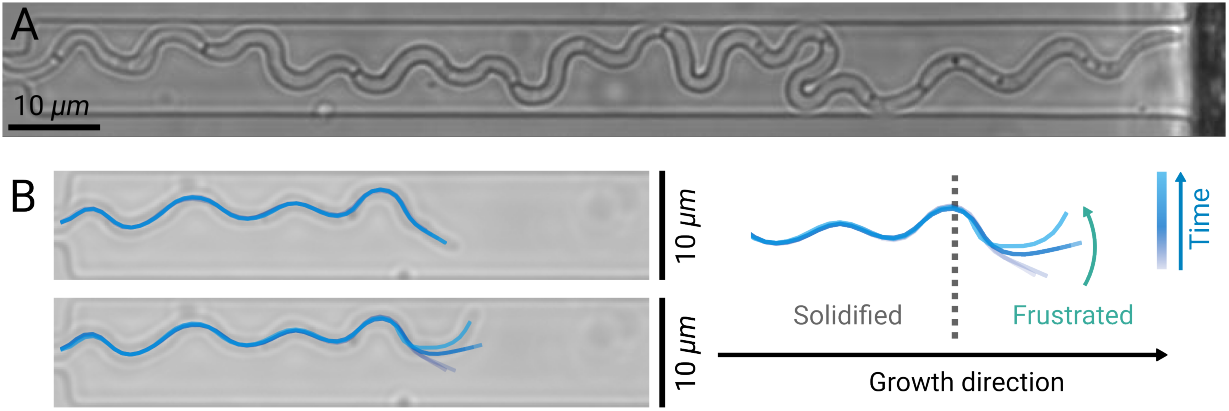
Curvature are built by successive sliding of an hyphal portion located at the apex. **A -** Image of an oscillating SC5314 *C. albicans* hypha within a 1.5µm high, 10µm wide microchannel. **B -** Images : Superimposition of the successive morphologies of the hypha shown in A at two different time points (blue lines). Schemes: selection of the 6 last successives morphologies highlighting the movement of the reconfigurable portion at the apex. **Fig.S3** - Caracteristic length of swiping. **Fig.S4** - Presence of the vacuole and solidification. **Movie 5a**: Timelapse of Fig.3a. **Movie 5b**: Tracking of timelapse of Fig.3a.

Any model of a growing hypha should now account for this new observation of the existence of a reconfigurable and a frozen part in a growing hypha. We propose such a model in the next sections, unifying our observations in both 2D and 3D within a single framework based on the concept of squeezed helices.

### The squeezed helix (squeelix) model of a growing hypha

Consider a helical Euler-Kirchhoff filament of length *L*, characterized by its arc length parameter *s*, Euler angles *ψ*(*s*),*ϕ*(*s*) and *θ*(*s*) describing the rotation of the material frame of the filament with respect to the laboratory frame axes [25]. Flattening the filament onto a plane constrains the third angle to *θ* = *π/*2 and its elastic energy is given by [21, 26, 1]

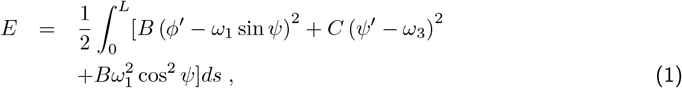

where *ϕ* (*s*) =: *κ*(*s*) stands for the filament curvature, *ψ*(*s*) designates the twist angle and *ψ* (*s*) =: *τ* (*s*) the twist, see also Fig. 4A. The two material constants *B* and *C* in Eq. (1) are the bending and torsional stiffness, respectively, *ω*_1_ and *ω*_3_ are the preferred curvature and twist of the unconfined three-dimensional helical filament [21]. Note that *ω*_1_ can be equated to the maximum curvature of the oscillations measured experimentally.

**Figure 4:**
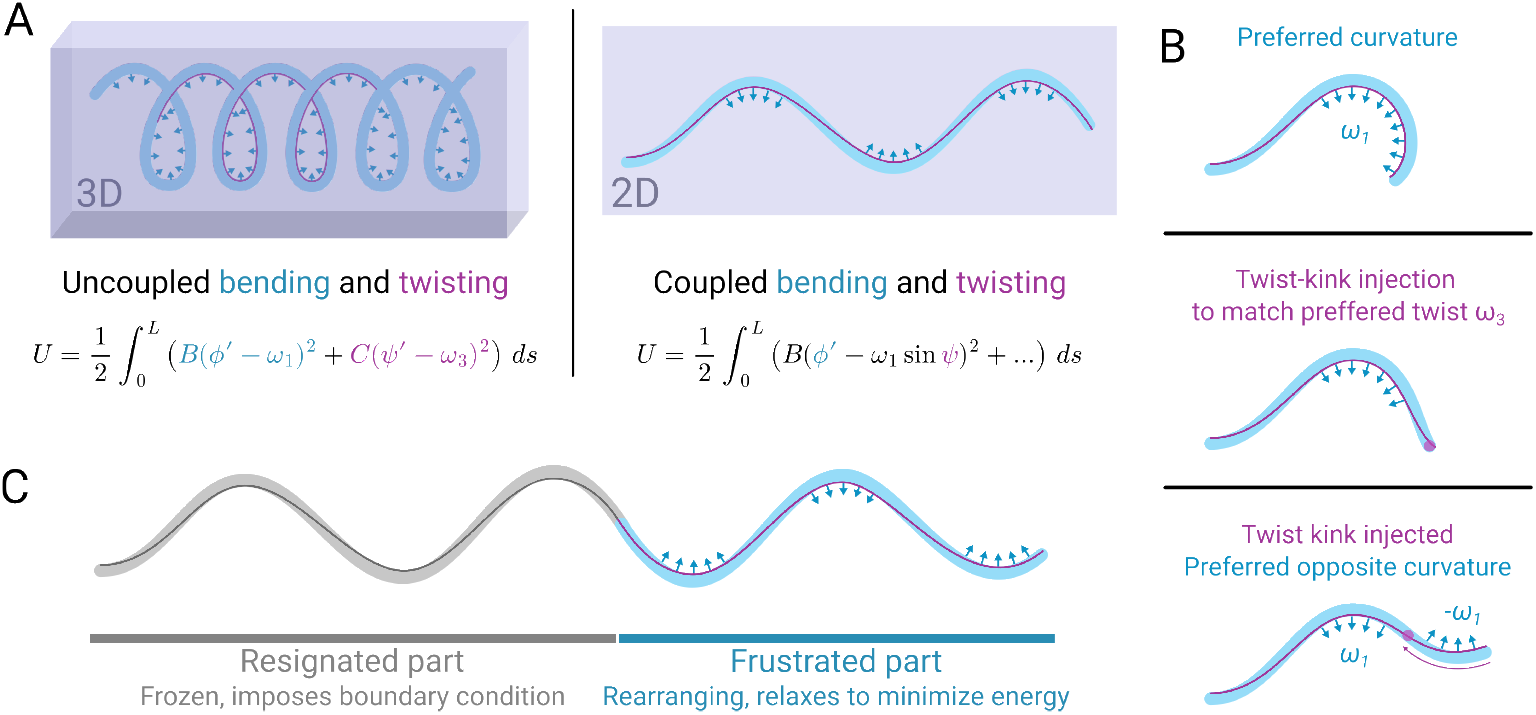
The “solidifying squeelix” model of a growing hypha. **A -** Illustration of the emergence of the 2D squeelix model by the confinement of a 3D elastic helical filament. Under confinement, the two Euler angles, *ψ*(*s*) and *ϕ*(*s*) couple with each other [21]. **B -** Successive shapes of a rearranging squeelix highlighting the occurrence of characteristic, discrete twist injection events (formation of twist-kinks) that correlate with sudden, intermittent, highly cooperative rearrangements characterized by curvature flipping within the tail portion of hyphae (“swiping”). **C -** Illustration of the solidification (“resignation”) process. A part at the tip is frustrated under the confinement, and described by the squeelix model, while the part further from the tip solidifies losing all the prestress with respect to the imposed shape in a process termed “resignation”.

The filament’s ground state resulting from minimizing the energy (Eq. (1)) obeys two coupled equations: the pendulum-like equation 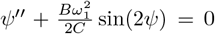 and (ii) *κ* = *ω*_1_ sin *ψ* indicating that curvature is slaved by the twist angle in contrast to the unconfined three-dimensional case (where both decouple). In general, depending on the material parameters, a rich variety of equilibrium shapes resembling loops, waves, spirals or circles exist [1, 26, 21]. These shapes can be seen as the result of interacting repulsive conformational defects corresponding to curvature inversion points. In terms of twist, such defects originate from a rapid variation of *ψ*(*s*) (reminiscent of a kink) and are called twist-kinks (Fig. 4B). A relevant dimensionless parameter is 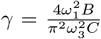 which measures the ratio of bending and twisting energy. For *γ >* 1, the ground state approaches a twist-kink-free circular arc of radius 1*/ω*_1_. For *γ <* 1, the filament can be populated by twist-kinks whose density is limited by their elastic repulsion energy [21].

#### Growth model

To model the growth of a *C. albicans* hypha, we recalled the observation that a filament is effectively separated into two parts: far from the growing tip, a solidified, shape-invariant part, and near to the tip, a dynamically rearranging part. An intuitive way understanding this division is to introduce the concept of stress-free, but curved reference shapes (far from tip) following the initial frustration of the tip portion/section and its subsequent rearrangement to find an energy minimizing, yet prestressed, position inside the plane. Both sections are markedly curved, yet only the tip proximal section is pre-stressed and has the potential to rearrange. The transition process from the frustrated to stress-relaxed state of the hyphae, we will refer to as “resignation” and the demarcation line between them the “resignation point”. The tip section progressively resigns with time, hardens, stops rearranging as it ages away from the tip. We simulated the combined growth-rearrangement-solidification process by minimizing the elastic energy of the rearranging part while progressively fixing (freezing) the shape of the filament along the resignation point, that is moving at same velocity as the growing tip. Each time step of the simulation consists of three iterative sub-steps: 1) A new short section of length Δ*s* in the reconfigurable portion is generated 2) The total elastic energy of this portion is minimized and 3) A section of same length Δ*s* is transferred from the reconfigurable to the solidified portion (Fig. 4C).

#### Validation and parameter matching

When scanning the range of parameters of a squeelix (preferred twist, curvature and the resignation length), we observe four different domains represented in the phase diagrams of **Fig. 5A-B**, established for three different lateral confinements provided by the channel (in plane) width *W*. Since the preferred experimental curvature (**Fig. 1D**) was close to 0.5µm^−1^, we chose this value to simulate the growth dynamics of hyphae confined in 10, 5, and 2.5 µm wide channels, with experimental validation for the smallest and largest widths. In a very satisfactory manner, the model captures the whole repertoire of hyphal growth behaviors observed under confinement. The swiping domain corresponds to hypha oscillations by sliding (**Fig. 5C**, see also **Supp Movie 5a**), already shown in Fig. 3 and **Supp Movie 6** for other examples. nosing (a term coined by Hanson et al.[27], describing the maintenance of a curvature at the growing apex) and looping (**Fig. 5D-F)** are occurring when the bending energy contribution dominates over twisting energy. Note that decreasing the channel width leads to a gradual shift towards large curvatures (small radii) of the looping domain. Experimentally, looping is the least represented behavior, as it requires the hypha to be able to go over another portion of itself, which is intrinsically limited by the z confinement. Nosing (**Fig.5E-F**, see also **Supp. Movie 7and 7b**), naturally emerges within our model. This kinetically trapped dynamic state results from the incentive of the hypha to change direction by generating a new twist kink, while not being able to do so due to the opposing force of the wall. The rearrangements due to the injection of every new twist kink require a certain minimal lateral (in plane) space to occur. The occurence of nosing thus increases as the width of the channel decreases, with a predominance for pure nosing at an intermediate width of 5 *µm*. At the swiping-nosing boundary, or for very thin channels, the hypha noses against a wall before switching to the other wall of the channel, creating bistable nosing. Such a behavior is also observed in our experiments (**Fig. 5F**, see also **Supp. Movie 8**) where the hypha slides for tens of microns on a single wall before switching sides.

**Figure 5:**
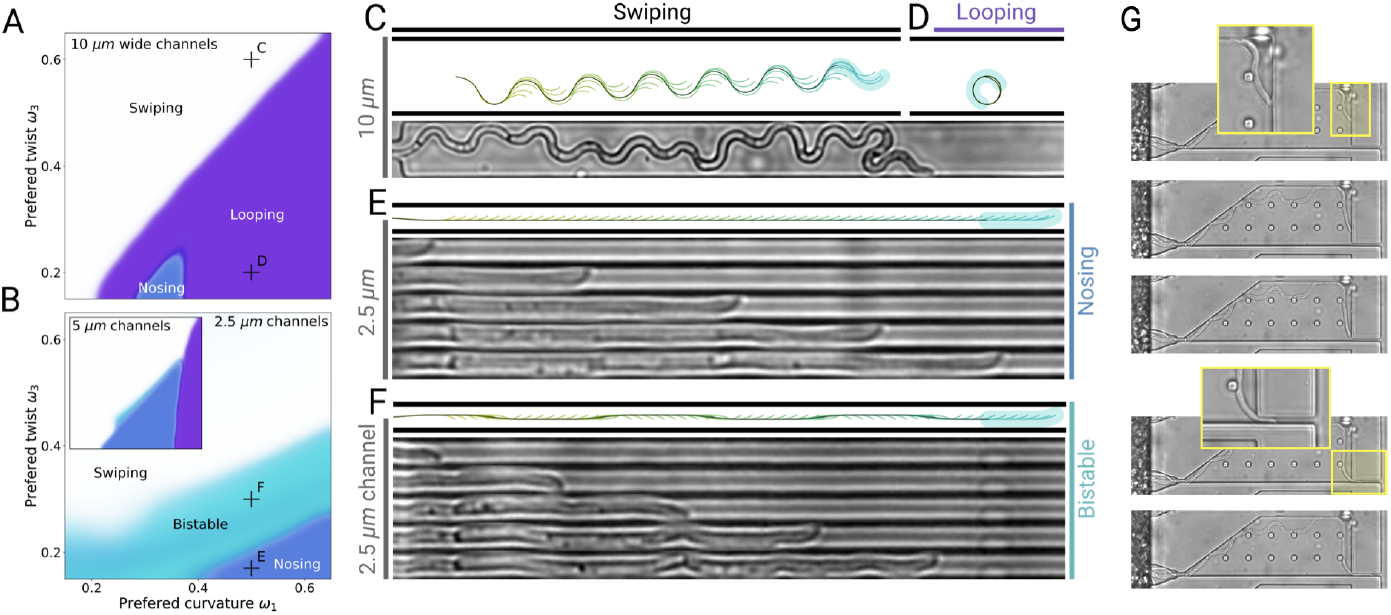
The squeelix model captures different observed behaviors of the hypha. **(A-B)** Phase diagrams established for a channel’s width of 10µm **(A)**, 5µm and 2.5µm **(B)**. Each phase diagram was computed using 18,000 growing squeelix. The resulting array was then smoothed and interpolated to obtain a continuous representation. **(C - F)** Simulation and comparison with experiments for swipping **(C)**, bistable nosing **(E)** and nosing **(F)**. These three squeelix modalities are supported by observations, contrarily to looping **(D)**. The shape at each few growth steps is displayed with a color, from yellow (old) to blue (new). The solidified part is represented in black, as well as the walls of the channels, and the hypha width is represented in light blue. For all simulations, the resignation length *L* and the preferred curvature *ω*_1_ are fixed with *L* = 8*µm* and *ω*_1_ = 0.5*µm*^−1^. The two other parameters are the channel width and the preferred twist (*W, ω*_3_) of the corresponding helical filament, whose values are respectively (10*µm*, 0.6*µm*^−1^), (10*µm*, 0.2*µm*^−1^), (2.5*µm*, 0.18*µm*^−1^) and (2.5*µm*, 0.3*µm*^−1^) for **(C), (D), (E)** and **(F). (G)** Nosing along walls and exit through a 3µm wide narrow channel. **Movie 6**: Larger view timelapse showing swipping in many channels. **Movie 7ab**: Timelapses showing nosing. **Movie 8**: Timelapse showing bistable nosing. **Movie 9**: Timelapse associated to Fig. 5G. **Movie 10a-d**: Simulations shown in Fig.5CD.

#### Relevance of nosing in path finding

We observed in large channels that nosing persists when the orientation of the walls change abruptly. Here, the regularly spaced pillars, whose function is to avoid the collapse of the microfluidic chamber, play a similar role as a wall in a thinner channel, by regularly confining laterally the nosing hypha. Strikingly, when the wall opens up a local, narrow gap - figuring a potential exit from the maze - the hypha engaging in nosing behavior manages to enter the gap and leave the channel (**Fig. 5G**, see also **Supp. Movie 8**). The maintenance of curvature at the tip near the point of contact with the wall (i.e. a quasi-perpendicular orientation of the tip tangent), allows the filament to find this opening without the need of any shape reconfiguration. This underlines the evolutionary advantage of nosing as a purely mechanical path finding algorithm, allowing to scan any continuity defect on a surface and be ready to invade the medium below through this weak point.

## Discussion

We have investigated the hidden, mechanically encoded algorithm behind the physical decision making process in growing *C. albicans* hyphae and show that they navigate their complex environments by turning their tip cells into helical sensory springs. Microfluidic tools were instrumental to control both the lateral and out of plane confinement of hyphae, allowing to fine-tune the level of environment-induced prestresses in the reconfigurable portion close to the hyphal tip. Hyphal growth within microchannels provided clear visual access to nosing behaviour and highlighted new classes of behavior such as bistable nosing, thus enabling the scanning of almost all morphologies associated with squeezed (as well as loosely confined) helices. Owing to 2D confinement-induced prestresses, the elastic energy stored in the reconfigurable portion of helices leads the tip cells to select the trajectory and ultimately the morphology of the filament as a whole.

The helical growth algorithm combines two important evolutionary features: i) improved scanning of the environment at the cellular scale, and ii) directional persistence of filament growth on super-cellular scales. The first phenomenon allows for the exploration and invasion of new habitats once they are reached, while in absence of any local environment change, the second phenomenon becomes more beneficial, by allowing the hypha to reach more favorable environments at minimal material and energy consumption cost - by taking the shortest path in space. This behavior is particularly reflected in the tangent correlation function of in-plane oscillatory hyphae. At short scales the tangent correlation function rapidly drops due to the “oscillatory snooping” environment exploration via the reconfigurable tip cell. However, the oscillatory snooping algorithm, being deterministically sinusoidal in nature, restores quickly the initial direction, leading to a recovery of directional persistence at larger distances.

One of the intriguing physical manifestations of the squeelix model is the prediction of nosing behavior [27], consisting in maintaining a curved tip during growth, and only appearing when the tip hits a wall and slides along it. This behavior has also been described in [28] in a situation of similar hyphal confinement as ours, and reported as a maintenance of the angle of approach in relation to the obstacle over time. Somehow, the nosing behavior appears to be a perfect strategy for an hypha to identify gaps along a wall, or weaknesses in a barrier in general, and use them as a gateway for invasion. It could be effectively utilized in the confinement situation provided by the 20-30µm internal diameter of intestinal crypts [29] or when navigating at the surface of any epithelia to implement paracellular route [30].

The pure nosing (in contrast to nosing-swiping) behavior is also a hint towards the presence of a simple form of memory in the growing hypha system. Indeed, owing to its binary decision to nose along the chosen wall (rather than the opposite one), once the channel is exited, the hyphae continue to move in a very similar direction, resuming upward or downward oscillation according to the wall (high or low) previously selected during nosing. The directional information has thus been stored over long times and distances in form of elastic tip deformation imprinted from one tip cell generation to the next one (see **Supp. Movie 11**: Nosing hyphae remember initial directions after leaving long, narrow channels).

Though at this point, the described phenomenon of (squeezed) helical growth and navigation, seems suggestive, we are yet to understand the biological processes behind picking the parameters of this helix under specific conditions. Estimates of the parameters, *ω*_1_ and *ω*_3_, that we can infer from fitting hypha shapes, show that they are themselves regulated and adaptable to the environmental conditions, like chamber height and even its lateral width.

An important experimental observation and central ingredient of our physical model is the existence of a moving frontier between an elastic, reconfigurable fixed length portion of filament at the tip, and a solidified growing part behind. Although such an abrupt transition remains a first order approximation, the fact that it provides a satisfactory description of our observations suggests the existence of a process of cell wall aging downstream the apex. Fungal cell walls are built from cross-linked polysaccharides (*β*-glucans and chitin) and are continuously remodeled through an equilibrium between synthesis and hydrolysis [31], mostly at their growing apex [32] but also all along the hyphal wall [33]. Although the changes in cell wall composition and architecture during aging have not yet been studied in hyphae, Silva et al. have recently reported in yeast cells of *Cryptococcus neoformans* and *Candida glabrata* a significant thickening of the cell wall in aged cells, accompanied with ultrastructural changes [34]. We cannot exclude that either cell wall thickening or increase of cross-linking occurs far downstream of the hyphal tip, as suggested in [32]. Closer to the tip up to a distance of about 15 µm, Chevalier et al. [35] reported in the filamentous fungus *Aspergillus nidulans* a monotonous decrease of the Young and surface modulus. This observation is in agreement with the ability of hyphae to bend near the apex to dynamically draw oscillatory growth. Interestingly, we observed that the reconfigurable portion is devoid of vacuole, and that the frontier between the frozen and reconfigurable portions is virtually determined by the front of the proximal vacuole (**Fig.S4**). Large vacuoles are associated to penetrative mechanical force in *C. albicans* through yet unknown mechanisms [36]. We might speculate here that they locally rigidify the hypha cell wall via overstretching by applying radial turgor forces.

Here we explored the macroscopic consequences of the helical scanning hypothesis and showed that many of the exploration and decision making events, like directional memory, oscillatory environment scanning and wall tracking, could be traced back to mechanical information processing encoded in the elastic energy minimization of a constrained and confined helix. This could be yet another natural example for the general phenomenon of “autonomous physical intelligence”, explored in manmade soft-robotics [37], where all the system responses to their complex environments are hard wired and coded within the subtle interplay of mechanical system parameters and geometry.

## Methods and Materials

### Fabrication of PDMS substrate and devices

PDMS thin films (experiments using agar gel): PDMS is spin-coated on a glass slide at 3000rpm, 30s, for an expected thickness of 30µm. The glass slide is then fixed with dental silicon at the bottom of perforated petri dishes, which were used as a culture chamber. The chamber can be exposed to an air plasma for typically a minute if an hydrophilic property is desirable before agar gel casting. Microfluidic devices: for the fabrication of the mold, a first layer of SU-8 2002 (MicroChem) is spin-coated on a silicon wafer to obtain a final thickness of either 1, 1.5 or 3µm. After a soft bake, this layer of negative photoresist is exposed to UV light thanks to direct laser writing lithography (Heidelberg, µPG101). Post-exposure bake and development with PGMEA are performed to obtain the first layer. Then a second layer of SU-8 2035 is spread out on top of the mold in order to get the macro-chambers of 30µm height. It is precisely aligned on the first level thanks to reference crosses on both layers. The mold is then exposed to UV light through a chromium mask (MJB4 mask aligner). The heights of the mold were measured with a mechanical profilometer (Dektak).

In an alternative version of this device, this first layer is itself a two-level layer with thicknesses of 1.5 and 6.5µm. The process of the 1.5µm layer fabrication is then repeated to add a second layer of 6.5µm height, an operation facilitated by the ease with which levels can be aligned with the µPG101.

The mold is first treated with trichloroperfluorooctylsilane (ABCR GmbH) deposited in vapor phase during 15 min in order to prevent adhesion of PDMS to the mold. Then PDMS is mixed with its curing agent (Sylgard 184 silicone elastomer kit) at the ratio 10:1 (w/w), degassed in bulk, poured on the mold, degassed again and finally cured at 75°C overnight. The cured PDMS is then peeled off from the mold, inlets and outlets are punched with a 1.5 mm diameter biopsy punch (Kai Industries). After the usual step of surface activation using an oxygen plasma, PDMS chips are bonded on glass Petri dishes (Fluorodish, VWR).

Overall, the first level corresponds to microchannels while a second, thicker level feeds these channels. It consists of an inlet opening onto a central channel extended by a serpentine and leading to the main outlet, and of two thin lateral chambers located on either side of the central channel connecting the microchannel outlet to the lateral outlets. As in [33], the serpentine was designed in order to equilibrate the flow toward the main outlet and the lateral outlet, allowing a few yeasts to be trapped in the pockets located at the entrance of each microchannel. These microchannel display different widths, from 1.5 to 40µm for further hyphal development.

### *Candida albicans* culture

Experiments were performed using the SC5314 reference strain of *C. albicans*. Strains are stored at −70°C in YPD medium (1% yeast extract, 2% peptone, and 2% glucose in distilled water (w/v)) supplemented with 25% of glycerol (v/v). They are streaked on YPD agar plates (YPD medium supplemented with 1.5% of agar (15 g/L of agar)) and kept for up to 3 weeks at 4°C. Prior to each experiment, a preculture is done: a colony from the plate is put in 3 *mL* of YPD liquid medium, and kept under agitation overnight at 30 °C. Typically, 50 µL of this solution diluted in 1mL of YPD corresponds to an optical density slightly below 1, corresponding to ≈10^7^ cells/mL. The 5 µL volume of this preculture (i.e. ≈10^6^ cells) is added to 495 µL of the filamentation medium composed of 445 µL of synthetic complete medium (i.e. SC medium, containing 0.67% yeast nitrogen base without amino-acids, 2% glucose, and 0.2% amino-acid mix (w/v)) and 50 µL of N-acetyl-D-glucosamine at 10mg/mL. Overall, the dilution of the preculture is made so that about a thousand of yeasts in the filamentation medium are inserted in the seeding chamber. To label the cell wall, we used calcofluor white staining at 1 mg/mL.

### *Candida albicans* seeding in agar gels

First, large quantities of agar are prepared and kept in their gel state at room temperature in closed containers to prevent evaporation. The gel is liquefied by microwaving and supplemented with N-acetyl-D-glucosamine at a final concentration of 1 mg/mL. The mixture is then cooled down to about 37°C to allow adding *C. albicans* yeasts without compromising their viability, and maintaining the gel in a liquid state. The gel + yeast solution is deposited on a surface consisting in a thin PDMS layer spin-coated on a glass slide at 3000 rpm for 30 s (for a final theoretical thickness of 30 µm), then cured for at least 2h at 75°C, and finally either treated by oxygen plasma (same protocol as to bond PDMS devices, see above) to make it hydrophilic or used right after curing.

### Optical imaging

Images and image sequences under bright-field illumination were obtained using an inverted microscope (Leica DMi8, equipped with a Hamamatsu C11440 camera) and Metamorph software, regulated at 37°C and 5% CO_2_ during time-lapse experiments (interval between each acquisition set to 5 minutes). We used a 40x dry objective (HC PC Fluotar 0.8NA) for multi-position acquisition. Fluorescent images were obtained using a Leica TCS SP8 confocal microscope (with LasX software) equipped with an oil-immersion 40x objective (APO 40x/1.30 oil CS2).

### Oscillation analysis (images)

In Fig. 2C the hyphae are observed on a free surface. To trace their shapes, we use the *Ridge Detection* [38] plugin for *Fiji* associated with a custom Python script to manually discard crossings between hyphae, and compute the tangent-tangent correlation functions. On the other hand, Fig. 1D hyphae are observed in microfluidic channels. Automatized gray-level image analysis is proving difficult due to the presence of the channel walls. Our approach consisted in subtracting an image of the channel before hypha growth, in order to eliminate the walls. In that aim, the position of the pre- and post-experiment images are precisely aligned using the *Template matching* plugin on Fiji to achieve a perfect overlap of the walls. The image is then enhanced on *Fiji* (using *background subtraction, dilate, remove outliers, contrast adjustment, image binarization*) to obtain a skeleton. After manual segmentation to discard crossing between hypha, *analyse particule* allows to select individual hyphae to be considered for final analysis. After a last step performed on *Fiji*, consisting at straightening hyphae (i.e. correcting any large-wavelength curvature), the amplitudes and wavelengths of each hypha are measured automatically using a custom-made Matlab program.

### Oscillation analysis (3D stacks)

A similar approach is used for 3D stacks. The stack is initially enhanced in *Fiji* using *contrast adjustment, Gaussian blur, thresholding, skeletonisation* and *dilation* to obtain a skeleton of the three-dimensional structure. Then, the *Simple Neurite Tracer* plugin [39] is used to semi-manually trace filaments. The tangent-tangent correlation functions are derived after interpolating and smoothing the traces in a separate Python script.

### Oscillation analysis (movies)

#### Preprocessing

To consistently track and add new points of a growing hypha across the frames of a time-lapse, another approach is necessary. The preprocessing is very similar, mainly consisting in stabilizing the movie using the plugin *Image Stabilizer* [40] from *Fiji* to ensure the hypha will be the only mobile object, applying a variance filter to enhance edges, and background substracting the first frame (with only the empty channels) from all other frames. This enhances the region of the movie where patterns have moved since the first frame, thus the growing hypha. The preprocessing ends by applying a Gaussian blur (with *σ* ≈ 1 *µm*) to remove the noise, and get a single blurred white line on a black background for the hypha.

#### Tracking

The hypha growth is tracked using a custom-made Python active contour algorithm: The filament is represented as a discretised curve of constant step, fully defined by its initial position, its total length, and its curvature at each point. Such a parametrized curve can be associated with a cost function that penalizes high curvatures and excess length, but rewards the part of the curves on the lighter pixels, corresponding to the hypha. Initially, we manually select the starting point of the hypha, its initial direction and length. This allows us to compute an initial guess for the first frame. At each frame, we optimize all curvatures to minimize the cost function using a gradient descent algorithm. This allows us to match the curve shape and length to the current shape and length of hypha. By iterating this process over the movie and using as initial guess for each frame the curve obtained from the previous frame, we can track along a movie the movements and the growth of the hypha.

### Squeelix model of a growing hypha

#### Squeelix model

The squeelix is modelled as a filament, living in a 2D plane, discretized by *N* points linked to each other by segments of fixed, constant length *ds*. The starting point is fixed, with a local orientation *ϕ*_0_, and a local preffered curvature direction *ψ*_0_. Each following point *n* is described by two varying parameters, the local curvature *ϕ*^′^_*n*_ and the local twist *ψ*^′^_*n*_. Initially, this set of parameters 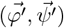 is set to arbitrary values, and associated with an energy, a discretized version of (eq 1). From there, the goal is to minimize the filament energy with respect to these parameters to find the optimal shape of the filament: the squeelix solution. This is done with the Sequential Least Squares Programming method (from *Scipy*.*optimize*.*minimize*). The choice of such method is justified by three reasons: it converges continuously to the nearest local minimum, making it closer to the fast, but continuous processes that drive the hypha rearrangement, with no large influx of energy to jump to far minimums of energy. It can also handle a few hundred parameters, as well as mathematical constraints on these parameters. Such constraints can be used to model which areas of the environment are accessible to the filaments and which are not, referring to PDMS walls in an experimental context. This is achieved using a scalar map of the plane where positive values indicate accessible areas and negative values indicate inaccessible areas. This allows us to model a large variety of lateral confinements.

#### Growth

In order to model the growth process, each time step is divided into two: the growth step and the rearrangement step. During the growth step, a new segment is added to the tip. At the resignation point, another segment hardens, effectively rebasing the frustrated filament on top of it while maintaining the original length. In the subsequent rearrangement step, the frustrated filament adjusts to find its optimal position. Overall, at each time step, the resignated part sets a new boundary condition, which influences the shape of the frustrated part after optimization. This optimized frustrated part then provides a new boundary condition for the resignated part, and the cycle continues iteratively.

#### Phase detection

To identify the phase (nosing, bistable, looping, or swiping) of a growing squeelix in a channel, we developed a Python detection method. First, nosing (and bistable) filaments are distinguished by evaluating the fraction of length the filament spends within a close range of a single (or both) walls of the channel. Then, discriminating looping filaments from swiping filaments is done by computing their aspect ratio.

## Acknowledgments

This work benefited from the technical contribution of the joint service unit CNRS UAR 3750. The authors would like to thank the engineers of this unit for their advice during the development of the experiments. This work was supported by the program “Helical Pathogen” ANR-23-CE45-0035, This project has received financial support from the CNRS through the MITI interdisciplinary programs, the Institut Pierre-Gilles de Gennes (“Investissements d’avenir” program ANR-10-IDEX-0001-02 PSL and ANR-10-LABX-31, and ANR-10-EQPX-34). Work in the laboratory of CdE is supported by the Agence Nationale de la Recherche (ANR-10-LABX-62-IBEID).

## Notes

### Competing Interest Statement

The authors have declared no competing interest.

