## Supplementary figures and images for "Snooping helices : The elastic path finding algorithm of growing hyphae"

### Fig. S1

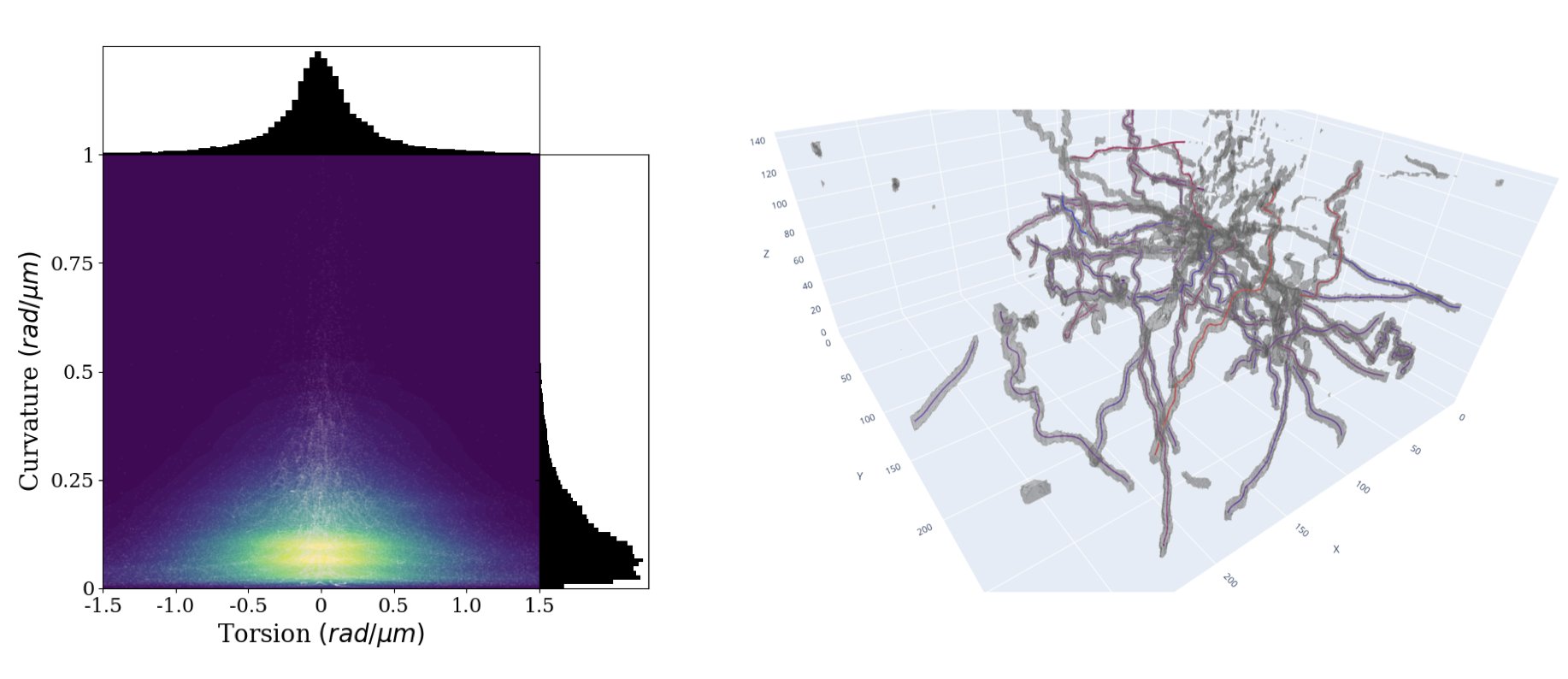

### Fig. S2

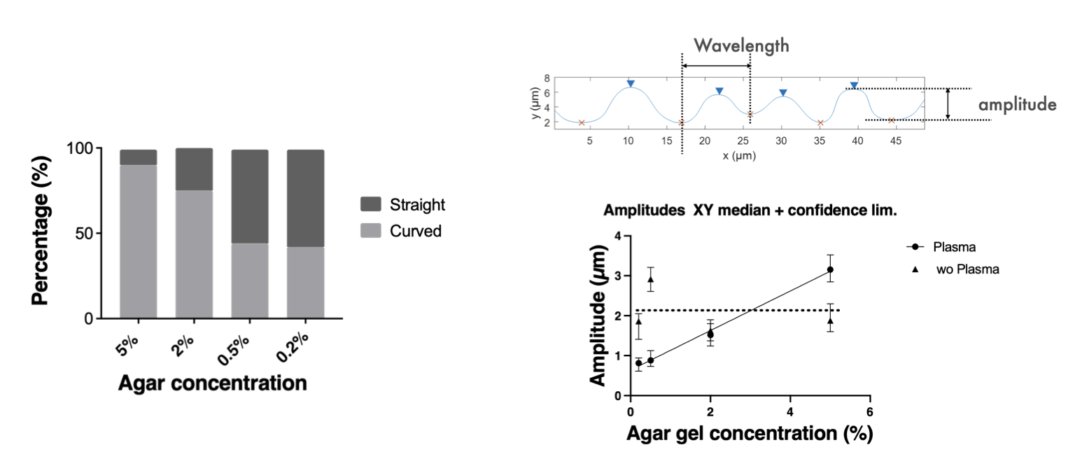

### Fig. S3

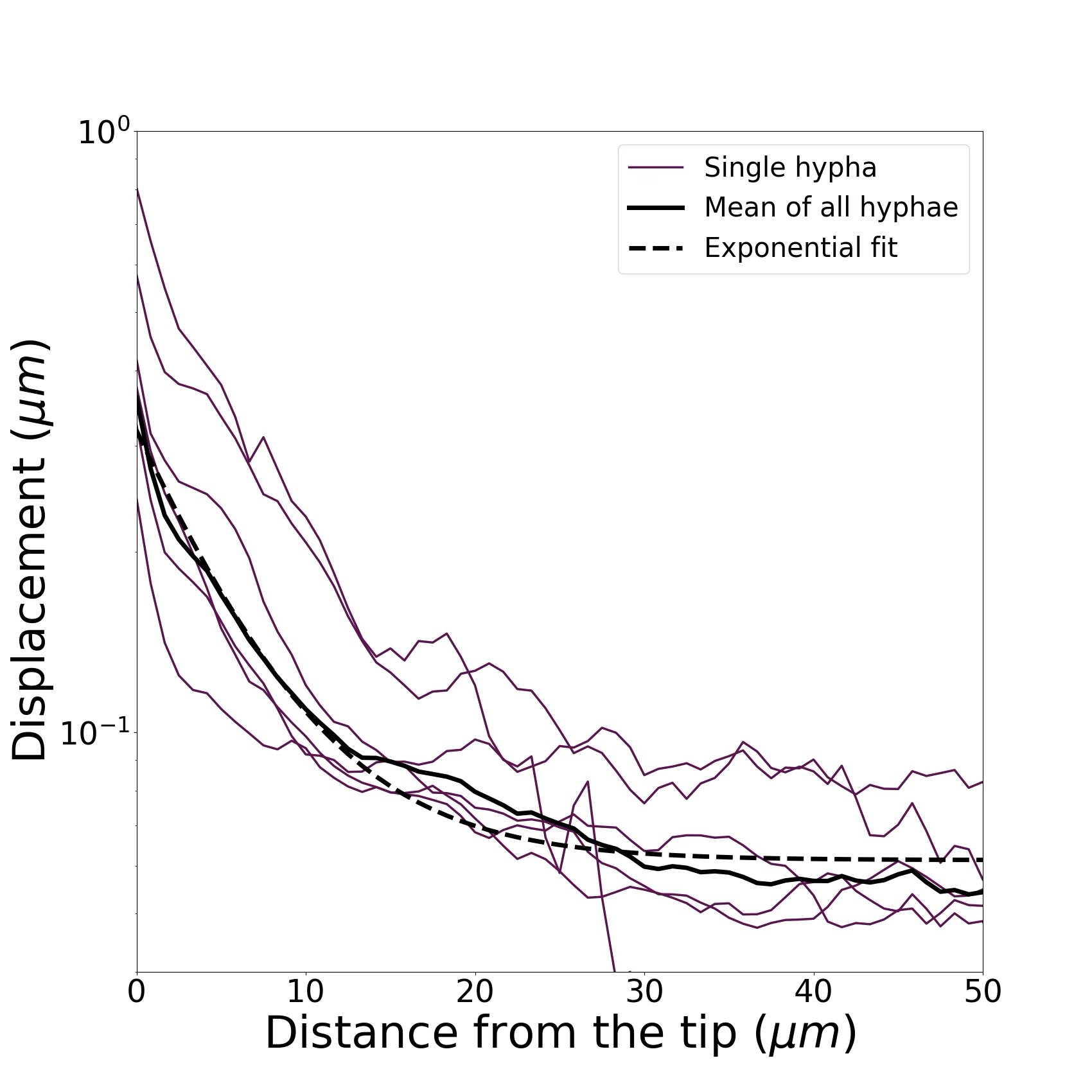

### Fig. S4

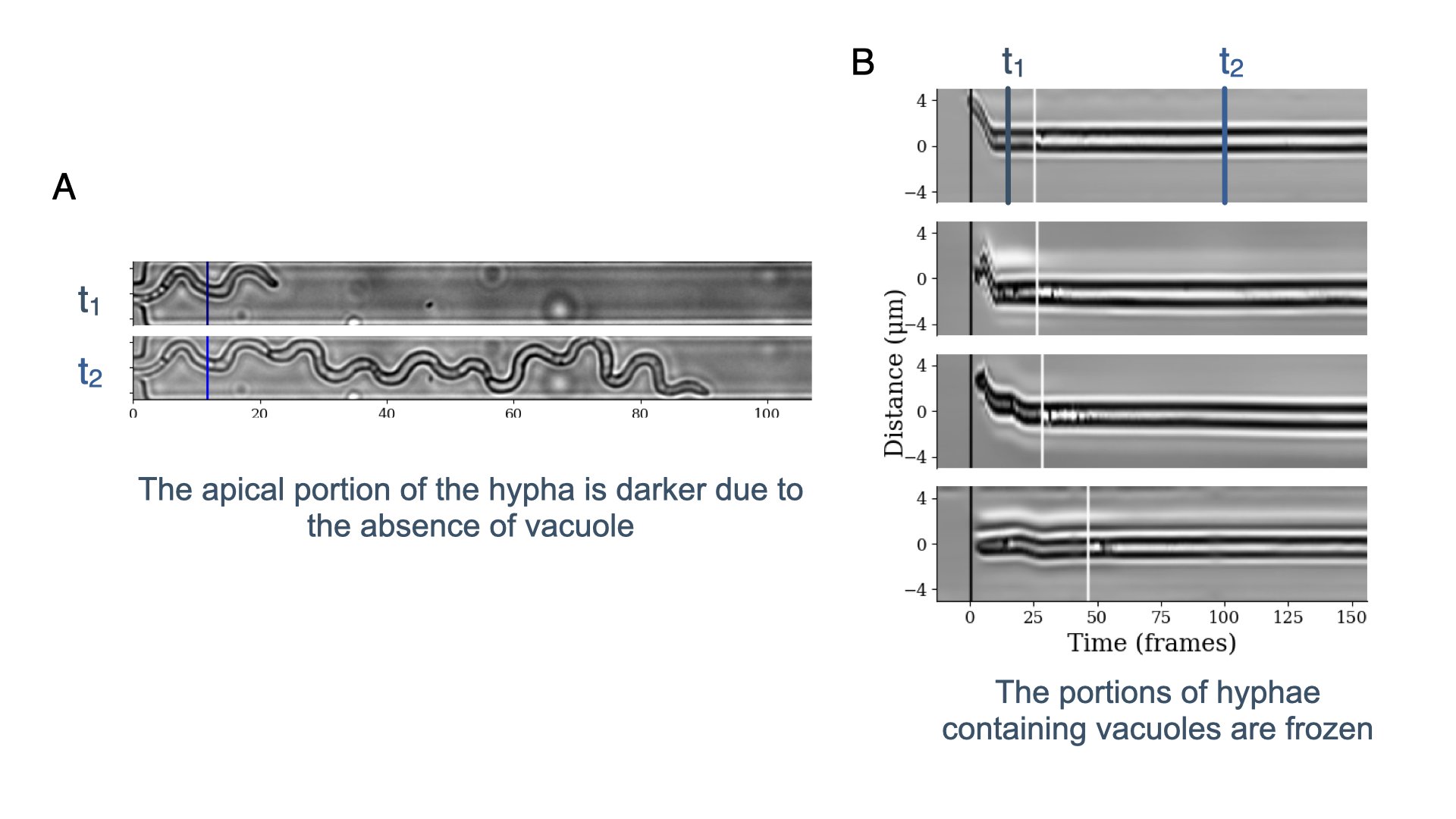
